# Identification of Novel miRNAs in Mouse Embryonic Stem Cells, Embryonic Fibroblasts, and Reprogrammed Pluripotent Cells

**DOI:** 10.1101/817395

**Authors:** Botao Zhao, Chunsun Fan

## Abstract

MicroRNAs (miRNAs) are a class of non-coding small RNAs that function in almost every known cellular activity. MiRNAs play an important role in gene regulation that controls embryonic stem cell (ESC) pluripotency and differentiation, as well as induced pluripotent stem cell (iPSC) reprogramming. In this study, we identified nine novel miRNAs by mining the deep sequencing dataset from mouse embryonic stem cells, mouse embryonic fibroblasts (MEF) and three kinds of reprogrammed pluripotent cells. Most of them are non-conserved but species-specific and cell-specific miRNAs. Eight miRNAs are derived from gene introns, including a “mirtron” miRNA, miR-novel-41. We also showed that miR-novel-27 is a mouse-specific miRNA and the 5′ arm of its precursor hairpin, embedding the mature miR-novel-27, uniquely exists in mouse species but not in any other *Placentalia* animals. Notably, the 5′ arm of the pre-miR-novel-27 hairpin shows nearly perfect palindrome to the 3′ arm suggesting that it was generated by inverted duplication of the 3′ arm. By this mechanism, the pre-miR-novel-27 hairpin was *de novo* gained in the mouse genome. This is a new type of *de novo* miRNA emergence mechanism in animals, which we called “inverted local half hairpin duplication” here. In addition, very limited nucleotide mutants accumulated on the newly emerged 5′ arm since its birth suggesting an especially young evolutionary history of the miR-novel-27 gene.

## Introduction

MiRNAs are endogenous, single-stranded small RNAs (∼22 nt) found in plants, animals and some viruses. They regulate target gene expression at the post-transcriptional level by binding to the complementary sequences within the target mRNAs. In animals, most miRNAs are transcribed from genome DNA sequences into large primary miRNAs (pri-miRNA) by RNA polymerase II [Lee et al., 2004]. The pri-miRNAs have hairpin structures that are then recognized and processed in the nucleus by the microprocessor complex, consisting of an RNA binding protein DiGeorge Syndrome Critical Region 8 (DGCR8 or “Pasha” in invertebrates) and a ribonuclease III enzyme, Drosha [Denli et al., 2004; Han et al., 2004]. The resulting hairpins are typically 60-100 nucleotides long and are called miRNA precursors (pre-miRNA) [Lee et al., 2003]. The pre-miRNAs are then exported to the cytoplasm by Exportin 5 (Exp5) and Ran-GTP complex [Kim, 2004; Lund et al., 2004; Yi et al., 2003]. In the cytoplasm, the pre-miRNAs are further processed into ∼22 nucleotides long miRNA duplex by the RNAse III enzyme Dicer [Grishok et al., 2001; Hutvagner et al., 2001]. Then, one strand of the duplexed miRNA is selected and incorporates into the RNA-induced silencing complex (RISC) to function miRNA-guided gene silencing [Iwasaki et al., 2010]. The selection of an active miRNA is primarily based on thermal stability. The strand with lower stability base-pairing of the 2–4 nt at the 5’ end preferentially associates with RISC and thus becomes the active miRNA [Frank et al., 2010; Khvorova et al., 2003; Schwarz et al., 2003; Suzuki et al., 2015]. The other strand, termed passenger miRNA or miRNA*, is favored for degradation and accumulates to a lower level than the active miRNA [Czech and Hannon, 2011; Yang et al., 2011]. Usually, the ration of mature miRNA/miRNA* is very asymmetric with a discrepancy of >10,000:1 [Liu et al., 2008]. Although the passenger miRNAs keep at a very low level, they still have a functional role in gene regulation [Mah et al., 2010].

To date, thousands of miRNAs have been identified in various organisms through cloning and sequencing or computational prediction. For example, the miRbase records 1,917 miRNAs and 1,234 miRNAs in human and mouse, respectively [Griffiths-Jones, 2004; Kozomara et al., 2019]. Although miRNAs were initially regarded as evolutionarily conserved, many non-conserved, species-specific miRNAs have been discovered particularly by deep sequencing [Bentwich et al., 2005; Zhang et al., 2018]. Most non-conserved miRNAs are expressed in specific cells at a low level and are believed to be evolutionarily young [Fromm et al., 2015; Ruby et al., 2007]. Interestingly, these young miRNAs are rapidly gained and most of them also rapidly lost indicating a rapid evolution of miRNA genes [Lu et al., 2008; Meunier et al., 2013]. In animals, the major source for miRNA birth is through *de novo* emergence of RNA hairpins from unstructured sequences, which occasionally acquired a promoter to enable successful transcription [Liu et al., 2008; Zhang et al., 2018]. For hairpins generated in introns, they have more chance to evolve to novel miRNA genes because they are innately transcribed along with their host genes and do not need the acquisition of transcription activity. Consistent with this evolution model, most newly identified novel miRNAs are from gene introns [Franca et al., 2017]. Some small introns can form pre-miRNA hairpins lacking the lower stem that can bypass the Drosha processing [Curtis et al., 2012]. These short intron derived miRNAs, called “mirtrons”, are confirmed in *Drosophila melanogaster* and *C. elegans*, and mammals [Babiarz et al., 2008; Berezikov et al., 2007; Curtis et al., 2012; Ladewig et al., 2012; Sibley et al., 2012]. Another source of miRNA emergence is local genomic DNA duplication of an existing miRNA gene to generate clustered miRNA gene family [Liu et al., 2008; Zhang et al., 2018]. However, the later miRNA emergence mechanism only explains the miRNA gene family expansion and evolvement but not the *de novo* birth of the first/progenitor miRNA gene of the family.

Initially, Ambros *et al*. established a set of criteria to annotate a miRNA, in which the expression of mature miRNA of ∼22 nt length is needed to be coupled with the existence of a stable hairpin structure precursor. Here, an ideal hairpin structure precursor must contain at least 16 complementary bases between the putative miRNA and the opposite arm. In addition, the mature miRNA should be phylogenetic conserved. Recently, Fromm *et al*. updated the miRNA annotation system [Fromm et al., 2015]. They stated that two 20–26 nt long reads from each of the two arms with 2-nt offsets should be detected and the expressed mature miRNA sequences should have 5′-end homogeneity. They further limited the loop size from 8 to 40 nt. The phylogenetic conservation criterion was not emphasized because many non-conserved miRNAs had already been validated as mentioned above. However, the ratio of mature miRNA/miRNA* is usually very asymmetric with a discrepancy of >10,000:1 [Liu et al., 2008]. It would be hard to detect miRNA reads from both arms especially if the miRNA expresses at a relatively low level. In this case, the expressed miRNA with no miRNA* read can still be considered as a miRNA candidate if they derive from an ideal hairpin structure [Axtell et al., 2011]. In this study, we applied the latest updated miRNA annotation criteria to successfully identified nine novel miRNAs in mouse ESCs, MEFs and three kinds of reprogrammed cells.

## Materials and Methods

### Cell samples and RNA data

The deep sequencing dataset was derived from our previous study and had been deposited in the Gene Expression Omnibus repository (www.ncbi.nlm.nih.gov/geo) with an accession number GSE52950 [Zhao et al., 2014]. Mouse embryonic fibroblasts (MEFs) and mouse embryonic stem cells (ESCs) (C57B6/129SvJae F1 strain) were cultured as described [Jiang et al., 2011]. Primary induced pluripotent cells (iPSCs) were established by induction of MEFs carrying dox-inducible four reprogramming factors (Oct4, Sox2, Klf4, and c-Myc). The nuclei of MEFs and iPSCs were transferred to enucleated oocytes to generate nuclear-transferred ESCs (NT-ESCs) and iPSCs (NT-iPSCs). Both ESCs and MEFs generated three replicate RNA samples and each reprogrammed pluripotent cell gave two replicate RNA samples. Total RNAs from these cell samples are subject to deep sequencing by the Solexa platform (BGI-Shenzhen, Shenzhen, China).

### Data analysis

After reads filtration and annotation [Zhao et al., 2014], unannotated small RNA were aligned to the Mouse Genome Assembly (GRCm38/mm10) and their surrounding genomic sequences were extracted for screening the existence of potential pre-miRNA-like hairpin structures by Mireap, a software developed by BGI-Shenzhen (http://sourceforge.net/projects/mireap/). After screening, a raw list containing 689 sequences was generated (Table S1). Because the median reads of all known miRNAs after normalization in these samples was 12.6 [Zhao et al., 2014], only small RNAs with normalized average reads more than 12.6 in at least one cell type were considered to be confidently expressed and chosen for this study. The normalization method and factors were the same as those used in our previous studies [Zhao et al., 2014; Zhu et al., 2019]. As the screening by Mireap was conducted just after the sequencing data was produced, some small RNAs in the raw list had already been identified to be miRNAs or other RNA types after that. Thus, further removal of these annotated small RNAs was performed. Then, the remaining sequences were folded by RNAstructure [Bellaousov et al., 2013] and the hairpin structures were manually checked according to the lastest miRNA annotation criteria [Ambros et al., 2003; Fromm et al., 2015]. In detail, a confident novel miRNA should range in size from 20 to 26 nt, with detectable expression and a “good” pre-miRNA hairpin structure [Fromm et al., 2015]. In this study, only potential small RNAs more than the median expression level of all known miRNAs were considered to be reliably expressed in these samples. A “good” pre-miRNA hairpin structure criterion used here has at least 16 -nt complementary base-pairing within the miRNA region and both arms separated by a single-stranded loop of 8-40 nt in length [Fromm et al., 2015]. The maximal free energy (MEF) threshold was set at −25 kcal/mol to ensure enough hairpin stability. In addition, the miRNA/miRNA* region should not be interrupted by unpaired bubbles larger than 4 nucleotides. The criterion that both arms should be detected was not forced to apply in this study. As mentioned above, the ration of mature miRNA/miRNA* is usually very asymmetric with a discrepancy of >10,000:1 [Liu et al., 2008]. Then, the detection of miRNAs from both arms will not be always expected especially if the miRNAs identified have much fewer reads than 10000. In addition, sequence conservation is omitted due to the fact that many non-conserved, species-specific miRNAs have already been identified and validated [Bentwich et al., 2005; Zhang et al., 2018]. Sequences conservation across species was viewed by UCSC Genome Browser (http://genome.ucsc.edu/) [Kent et al., 2002] and alignment was performed by Clustal X [Jeanmougin et al., 1998]. All long RNAs were folded and presented by RNAfold web server [Gruber et al., 2008].

## Results

To identify potential candidate miRNAs, unannotated small RNAs were aligned to the Mouse Genome Assembly (GRCm38/mm10). Their surrounding genomic sequences were extracted and folded as RNA sequences for the potential existence of pre-miRNA-like hairpin structures. 689 potential hairpin structures were identified (Figure S1). Among them, only small RNAs with average normalized reads more than 12.6 (the median reads of all known miRNAs in these samples) in at least one cell type were considered to have confident evidence of expression. After further removal newly identified miRNAs and other small RNAs overlapped with known snoRNAs, exon of protein-coding genes, and so on, a list containing 63 small RNAs was generated. Their potential pre-miRNA hairpin structures were manually checked to ensure to satisfy with the recently updated miRNA annotation criteria [Fromm et al., 2015]. Finally, nine candidates of pre-miRNA hairpins were identified (Table 1) and their structures were presented in Figure 1. All these hairpins are reliably stable with high fold possibility and low folding free energies (Figure 1& Figure S1). Remarkably, both arms of pre-mir-novel-27 and pre-miR-novel-65 produced mature miRNA sequences (Table 1). The 2-nt 3′overhang between the two miRNAs and their miRNA* sequences indicated a typical Drosha/Dicer processing character (Figure 1). Taken together, all the evidence strongly supported that these hairpins are *bona fide* precursors of miRNAs. Consistent with the previous evolution model, seven of the nine miRNA hairpins derive from the gene introns indicating a high ratio of intron derived miRNAs (Table 1) [Franca et al., 2017].

**Table 1.**
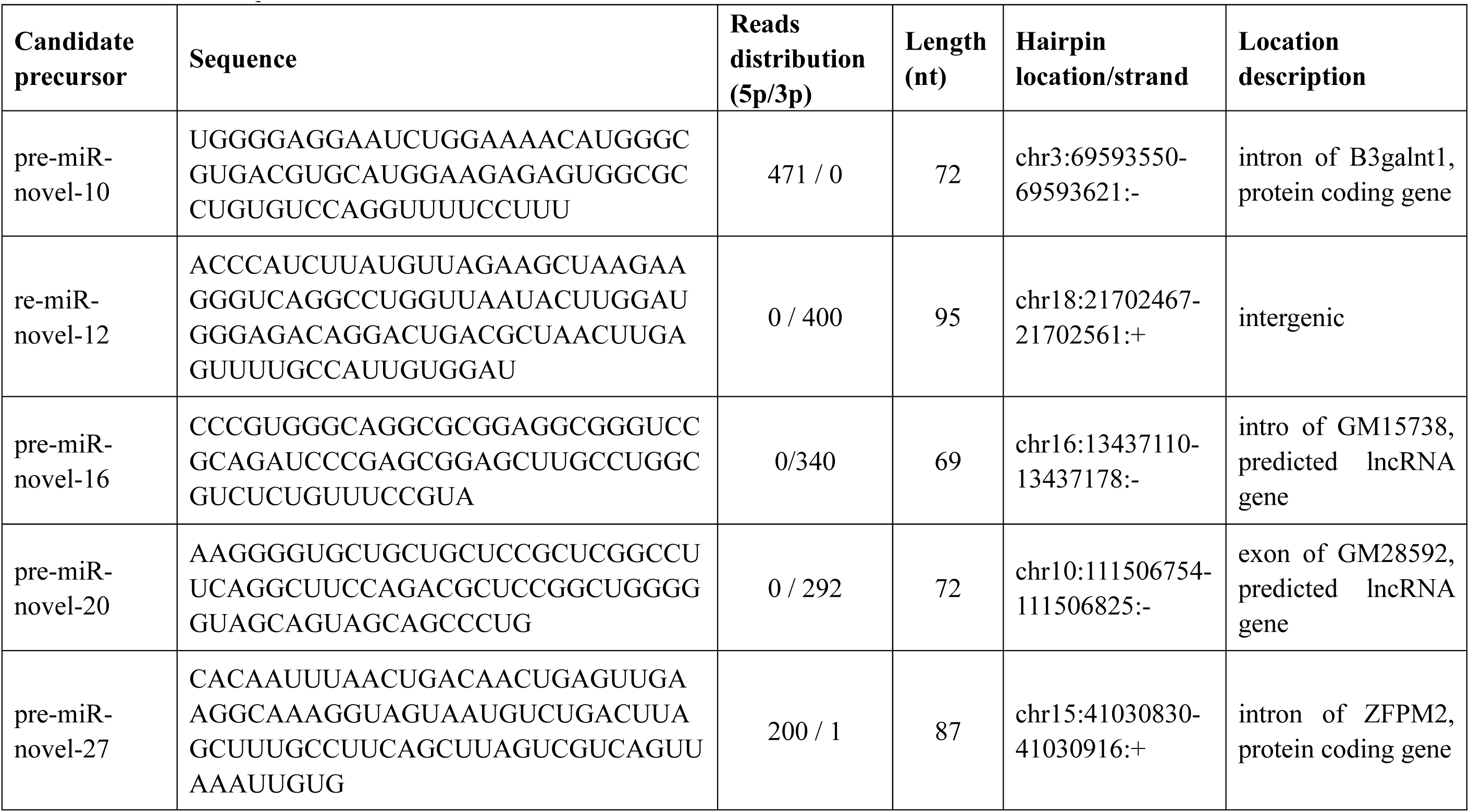
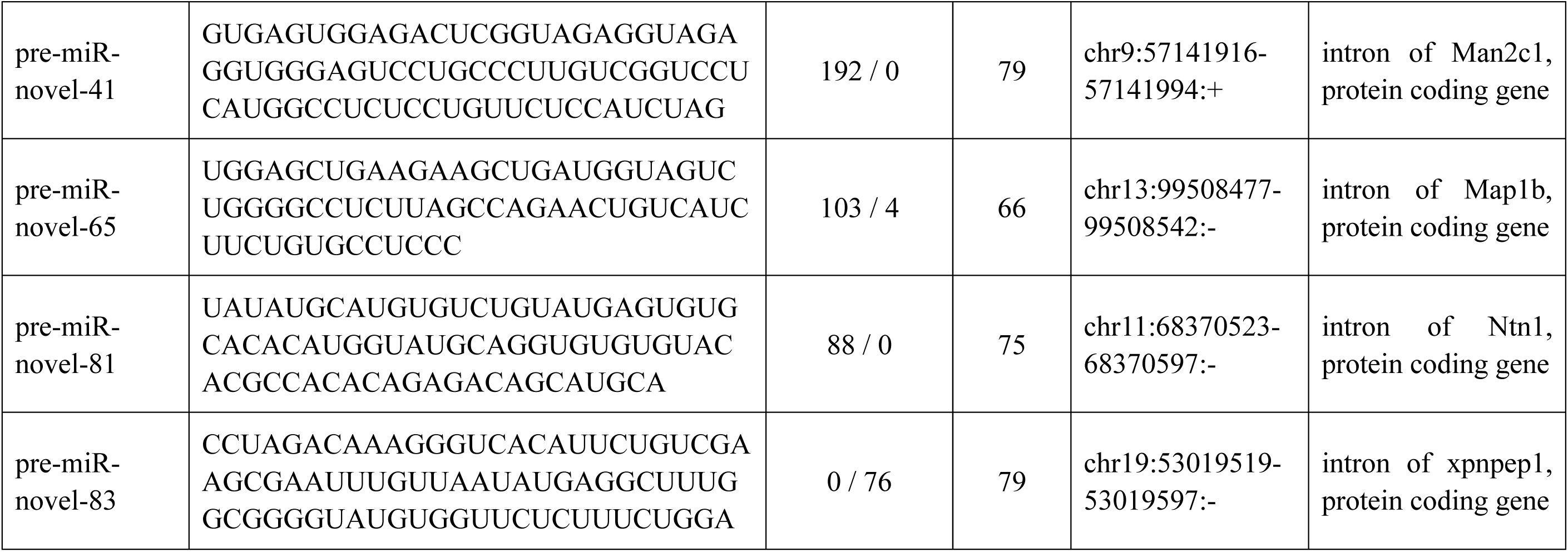
The hairpin sequences of the identified candidate miRNAs.

**Figure 1.**
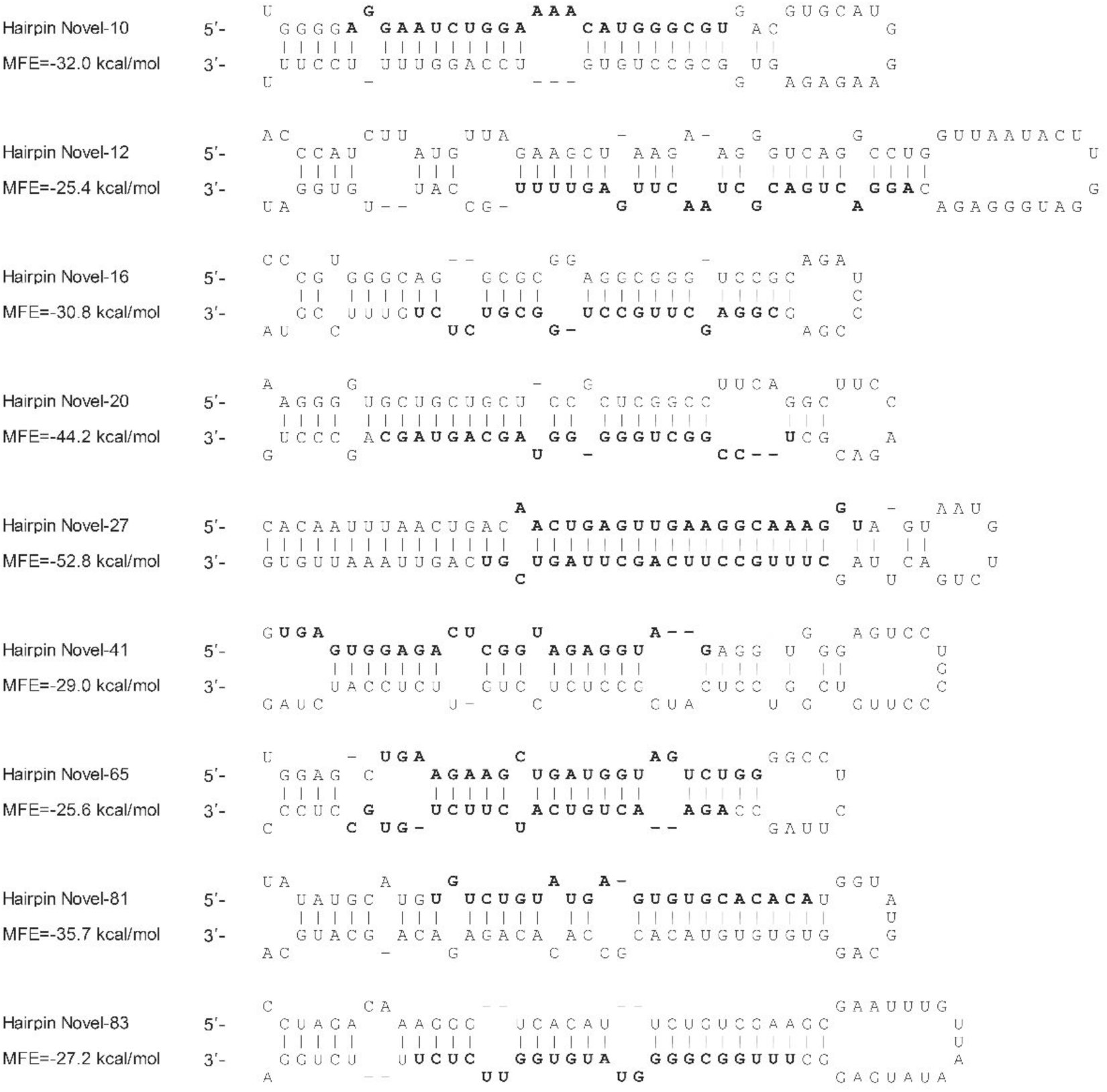
Hairpin structures of the candidate miRNA precursors were shown as text format. Detected mature miRNA sequences were indicated as bold nucleotides. Their reads were listed in Table 1. Maximal Free Energies (MFE) of the hairpins were listed at left.

Then, we searched these small RNAs against the initial 689 small RNAs to include their diverse sequences with different 5′- or 3′-ends. Four miRNAs, miR-novel-12, miR-novel-20, miR-novel-41 and miR-novel-81, had diverse reads with different 3′-ends, and yet, all the diverse reads of each miRNA had 100% 5′-end homogeneity (Table 2). After combining these diverse reads, miR-novel-10 showed the highest expression with 471 total reads in all samples, followed by miR-novel-12 with 400 total reads and miR-novel-16 with 340 total reads, respectively. Interestingly, most of these miRNAs dominantly expressed with more than half reads in one cell type (Table 2 & Figure 2). Four novel miRNAs (miR-novel-10, miR-novel-20, miR-novel-41 and miR-novel-83) were dominantly expressed in mouse ESCs. MiR-novel-12, miR-novel-16, and mir-novel-81 showed high abundance in iPSCs. MiR-novel-27 and miR-novel-65 mostly expressed in MEFs and NT-ESCs, respectively (Table 2 & Figure 2). The normalized reads in all samples were listed in Table S2.

**Table 2.**
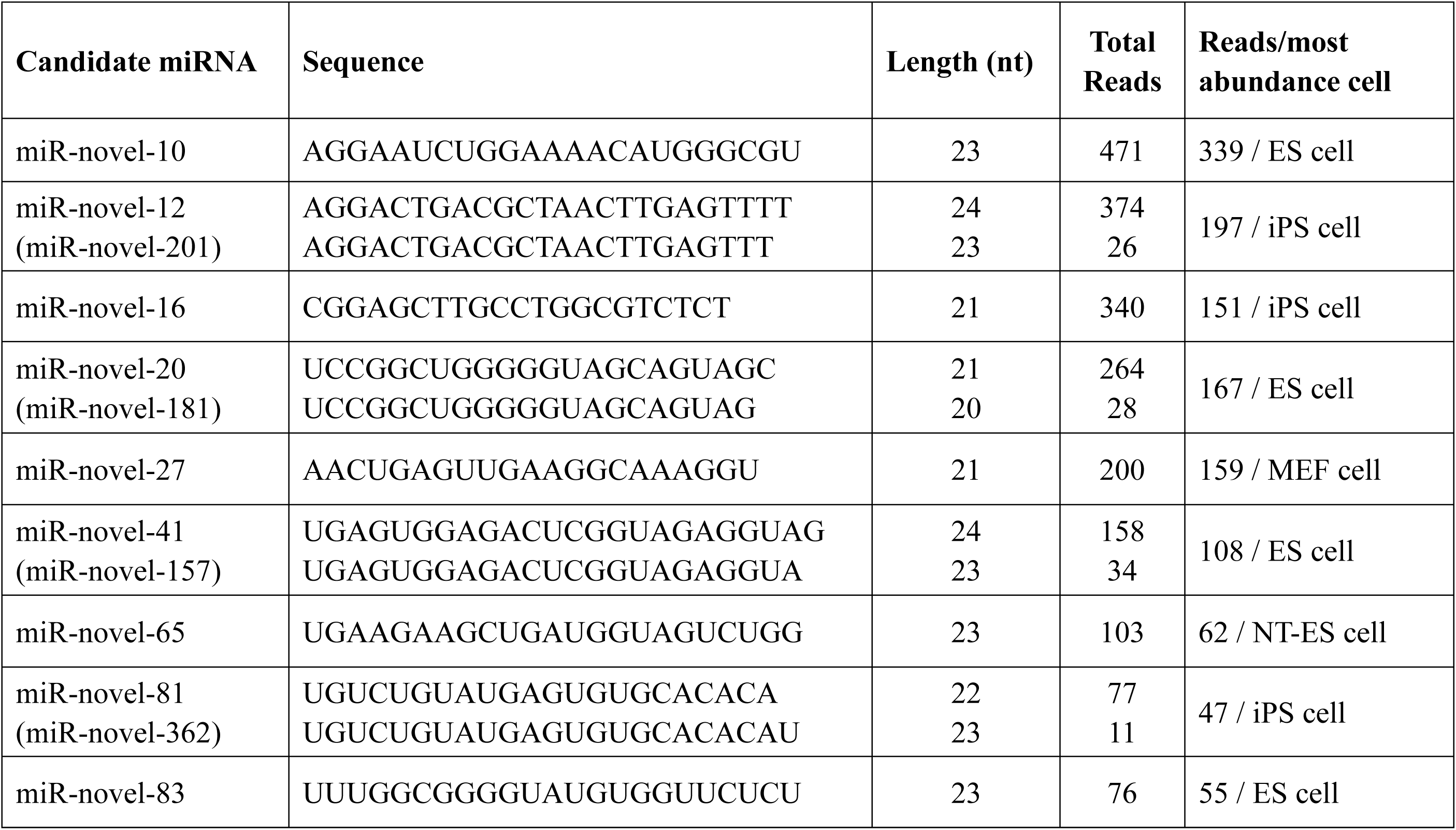
The mature sequences of the identified miRNAs.

**Figure 2.**
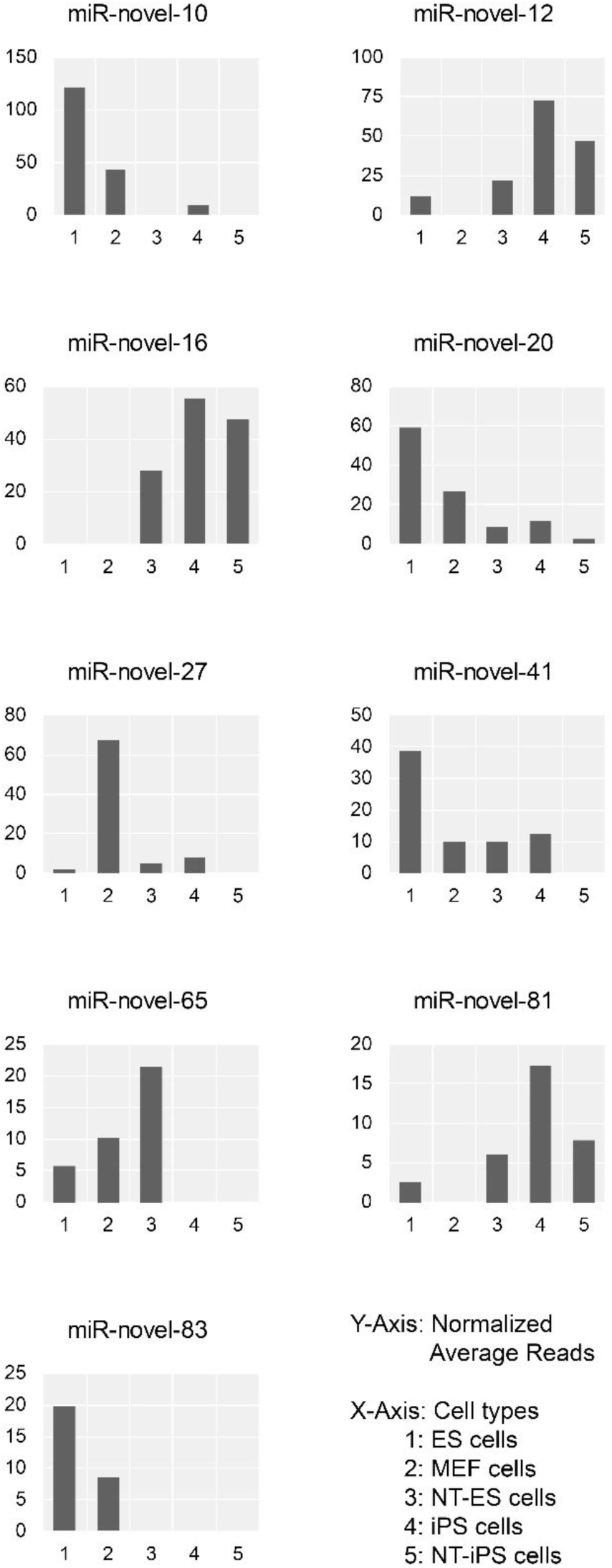
Normalized average reads of the candidate miRNAs in each cell type. ES cells: mouse embryonic stem cells (three replicate samples); MEF cells: mouse embryonic fibroblasts (three replicate samples); NT-ES cells: nuclear-transferred mouse embryonic stem cells (two replicate samples); iPS cells: induced pluripotent cells (two replicate samples); NT-iPS cells: nuclear-transferred induced pluripotent cells (two replicate samples).

As mentioned in Table 1, miR-novel-20 overlaps with the exon of GM28592, a predicted lncRNA gene. To date, there is no functional information reported about this lncRNA. Very limited data in Expression Atlas shows that GM28592 expresses at a low level in the mouse brain, liver, and epidermal ectoderm at embryonic day 9.5 (https://www.ebi.ac.uk/gxa). To avoid mis-annotation, we checked that if the hairpin structure of pre-miR-novel-20 could be stably formed in the context of the full-length RNA of GM28592. As shown in Figure 3, the full-length of GM28592 RNA can form the intact pre-miR-novel-20 hairpin with a very high probability that could serve as a substrate of the miRNA microprocessor complex. Combining with the miR-novel-20 reads detected in this study, we concluded that GM28592 is more likely a miRNA gene although it may also function as a lncRNA.

**Figure 3.**
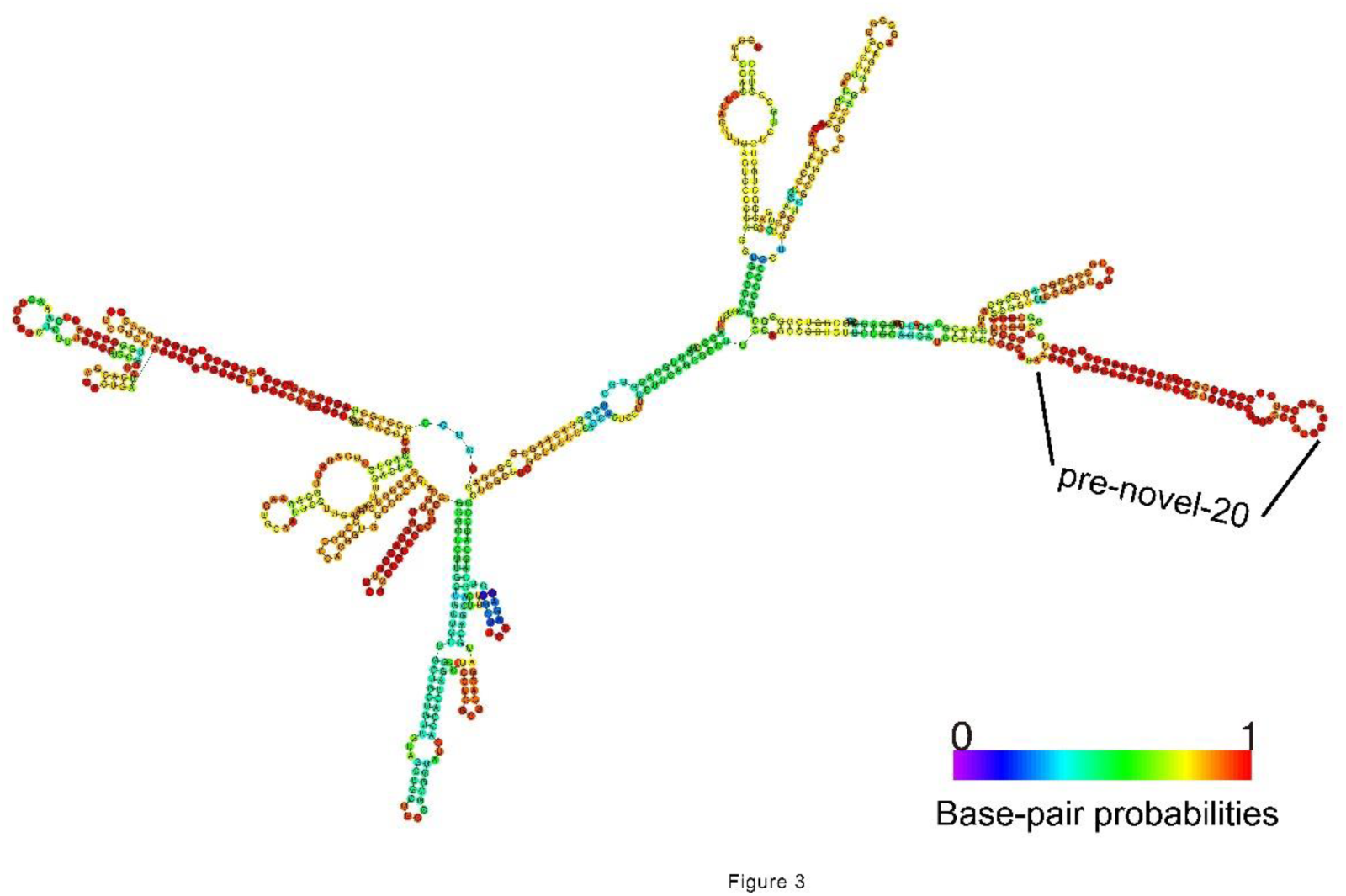
The folded full-length RNA of predicted GM28592 gene. The location of the pre-miR-novel-20 was indicated. The colors of nucleotides showed the base-pairing probability as the scale bar.

MiR-novel-41 derives from the intron of Man2c1, a protein-coding gene (Table 1). The host intron is 79-nt and could perfectly fold into a pre-miRNA hairpin structure from which miR-novel-41 is produced (Figure 4). The formed hairpin structure lacks the lower stem part, a typical feature for bypassing the Drosha processing (Figure 4). Mammalian mirtrons predominantly produce mature miRNAs from the 5′ arm. In these cases, the corresponding 3′ region is particularly pyrimidine-rich. Consistently, miR-novel-41 also comes from the 5′ arm and there is a very high ration of C and U throughout the 3′ arm, especially in the miR-novel-41 complementary region (Figure 4). With these typical characteristics, miR-novel-41 was confidently considered as a mirtron miRNA here. In addition, miR-novel-41 starts at the second nucleotide on the 5′ arm indicating it is a 5′ tailed-mirtron [Curtis et al., 2012].

**Figure 4.**
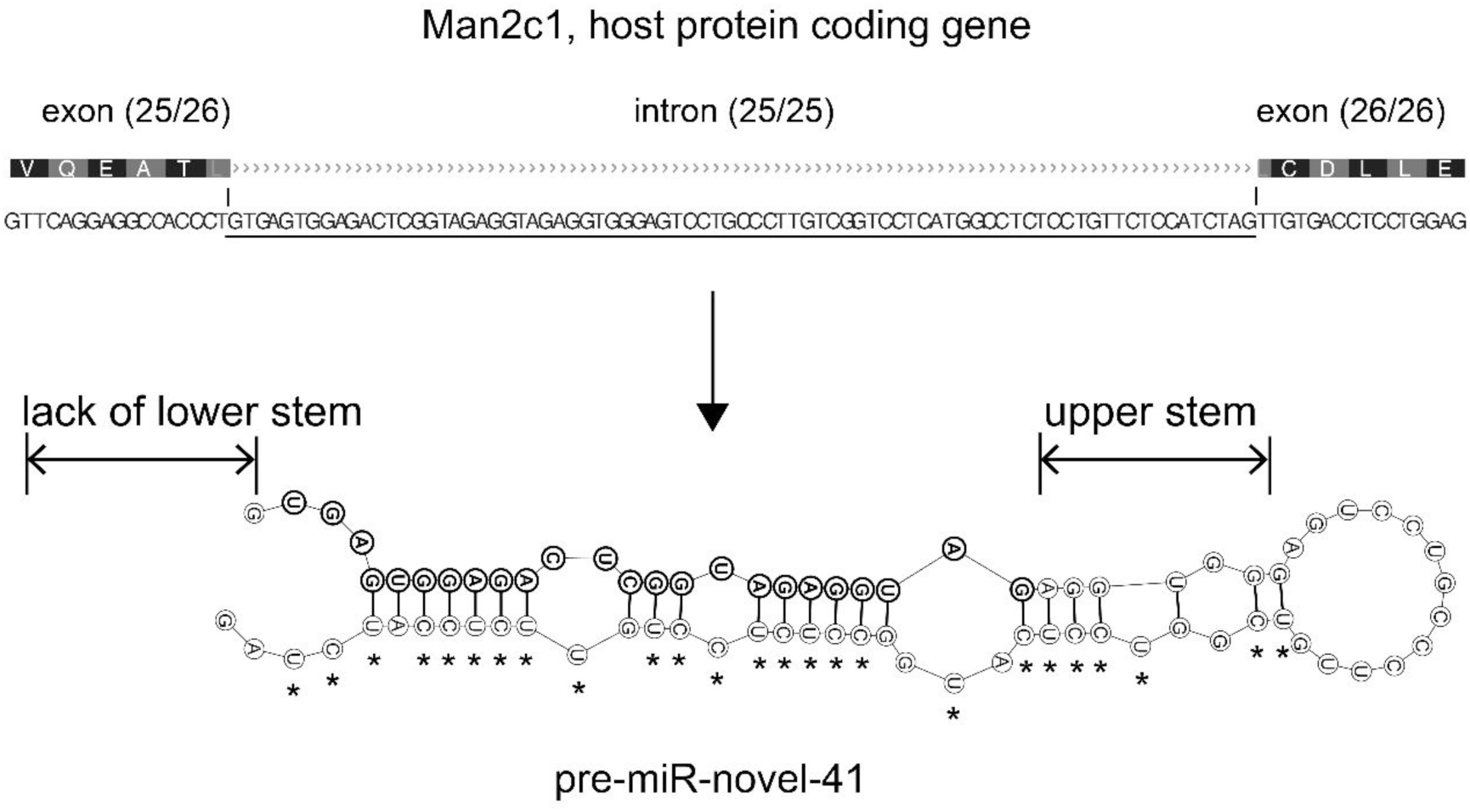
MiR-novel-41 is a mirtron miRNA. The sequence of the 25th intron of Man2c1 was underlined. The folded hairpin structure was lack of lower stem part as shown. The mature miRNA-novel-41 sequence was labeled by bold fonts and the pyrimidine nucleotides on the 3′ arm were indicated by asterisks.

It has been extensively validated that there are many non-conserved, species-specific miRNAs [Bentwich et al., 2005; Zhang et al., 2018]. To check the sequence conservation of novel miRNAs identified in this study, the pre-miRNA sequences were aligned across *Placentalia* animals by UCSC Genome Browser (http://genome.ucsc.edu/) [Kent et al., 2002]. Pre-miR-novel-20 and pre-miR-novel-41, especially their seed sequences, show well conservation in most *Placentalia* animals. For pre-miR-novel-65 and pre-miR-novel-83, although they show somewhat conservation in some *Euarchontoglires* animals, their mature miRNAs are not conserved. Four miRNA precursors, pre-miR-novel-10, pre-miR-novel-12, pre-miR-novel-16, and pre-miR-novel-81, only show high conservation within *Murinae* animals, so do their mature miRNAs. Interestingly, miR-novel-27 is a mouse-specific miRNA and does not exist even in rat species. Then we narrowed down the species to *Murinae* animals, which only include mouse and rat in the UCSC Genome Browser database. As shown in Table 3 and Figure S2, all these novel miRNAs, except for miR-novel-27, are well conserved in mouse and rat. For the seed region, five miRNAs (miR-novel-12, miR-novel-16, miR-novel-41, miR-novel-65, and miR-novel-81) are 100% conserved between mouse and rat (Table 3 & Figure S2). Mouse miR-novel-10, miR-novel-16 and miR-novel-85 have 2, 2 and 1 nucleotides different from their rat paralogs, respectively (Table 3 & Figure S2). Four of the five different nucleotides are G/A or C/T substitutions and miRNA seed sequences bearing such type of nucleotides substitution could still bind to the same target sequences because the G:U base-pairing is allowed in miRNA/target binding (Table 3 & Figure S2). Overall, all these novel miRNAs, except for the mouse-specific miR-novel 27, have conserved miRNA/target gene recognition relationship.

**Table 3.**
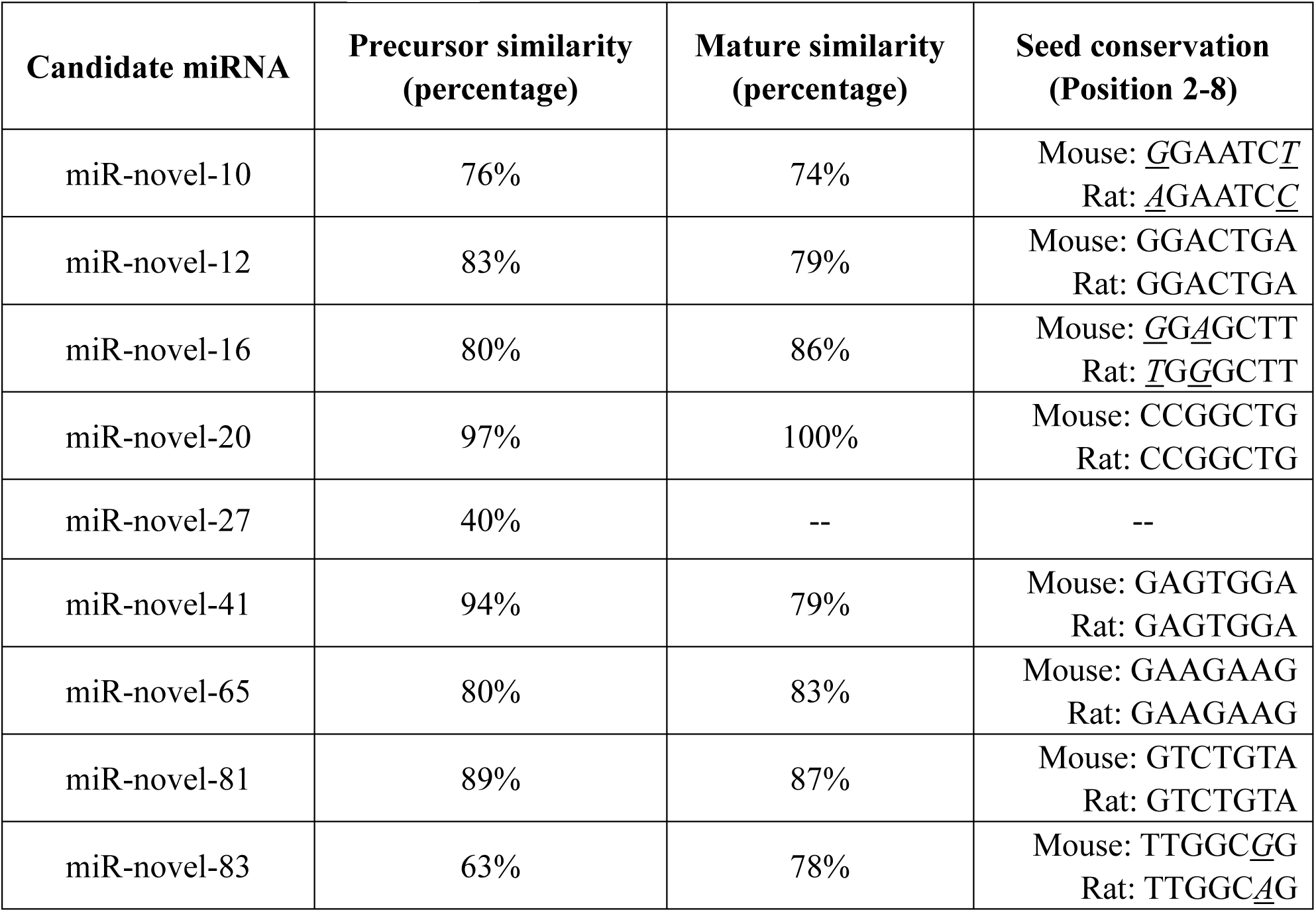
The sequence similarities of the miRNAs between mouse and rat species. Different nucleotides in the seed region between the two species were indicated by underlined italic fonts.

For pre-miR-novel-27, it is very strange that its 5′ arm embedding mature miR-novel-27 is unique to mouse species and does not exist in all other *Placentalia* animals; while the host intron of ZFPM2 gene is highly conserved across these animals (Figure S3). Therefore, RNAs from this location of any *Placentalia* animals but mouse lack of the 5′ arm sequence and could not form the hairpin structure of pre-miRNA-27 at all (Figure S4). It is not possible that all other animals lost this short sequence but the mouse kept it during evolution. Instead, it is obvious that the mouse genome gained this sequence and this gain event must occur after the split of mouse and rat species. This indicates that miR-novel-27 should be a very young miRNA. To explore how miR-novel-27 originated, we looked close to its precursor hairpin structure and sequence. The stem part of pre-miR-novel-27 shows nearly perfect complementary base-pairing that is much higher than most known miRNAs (Figure 1). Such high base-pairing indicates the existence of a palindromic sequence on its genomic DNA and also suggesting that the *de novo* emergence of the 5′ arm came from a short inverted duplication of the 3′ arm (Figure 5A). RNA from such a palindromic DNA sequence can natively fold as a hairpin structure with a full complementary stem part. As a result, the 5′ arm of the pre-miRNA hairpin was *de novo* gained. This is a new type of miRNA gene *de novo* birth mechanism, never revealed before in animals, with miR-novel-27 as a supporting example. This mechanism only duplicates the half hairpin and is distinct from the previously demonstrated local genomic DNA duplication of the full sequence of an existing miRNA gene, the major source of miRNA gene family expansion. We called this new miRNA birth mechanism as “inverted local half hairpin duplication” in order to distinguish from the previously demonstrated local duplication of existing miRNA genes (full hairpin). Here, we presented the new type of *de novo* birth and evolution model of the miR-novel-27 gene in Figure 6.

**Figure 5.**
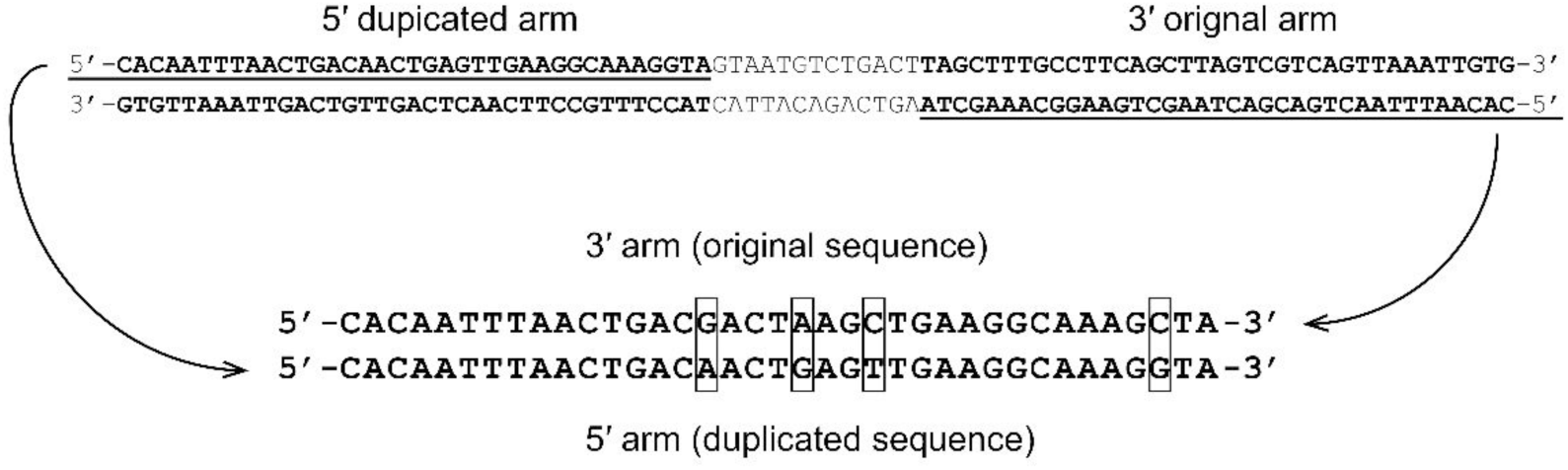
The palindromic sequence of the pre-miR-novel-27 hairpin. The 3′ original arm and 5′ duplicated arm were back-to-back compared and the different nucleotides were boxed out.

**Figure 6.**
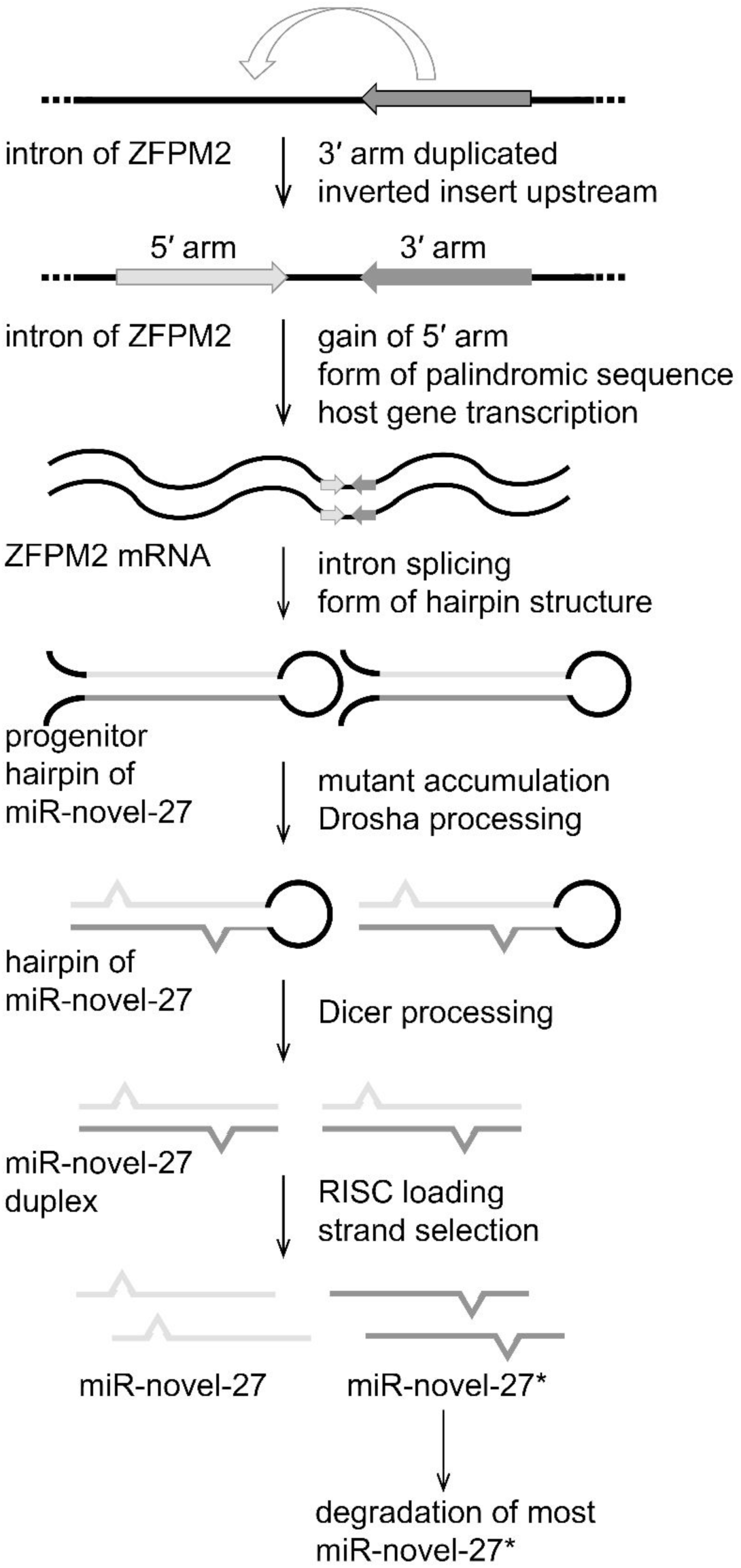
The inverted local half hairpin duplication model explained the *de novo* birth and evolution of the miR-novel-27 gene. The gray sequence indicated the original 3′ arm/miR-novel-27* and the light gray sequence indicated the duplicated 5′ arm/miR-novel-27.

Since pre-miR-novel-27 hairpin emergence, mutants gradually accumulated with the course of evolution, and the hairpin eventually was recognized by the miRNA microprocessor complex and produced the miR-novel-27 duplex (both strands detected in this study) with 2-nt 3′ offset. Today, there are only four different nucleotides between the original 3′ arm and the duplicated 5′ arm of the pre-miR-novel-27 hairpin, and therefore, the base-paring of the stem part is much higher than most of the known miRNAs (Figure 1). Considering the mutation accumulations on each arm are roughly equal in theory, we could expect that only two nucleotides mutations accumulated on the newly emerged 5′ arm since its birth. This information suggests that this inverted local half hairpin duplication event might occur very recently and further supports that the mouse-specific miR-novel-27 is an especially young miRNA gene.

## Discussion

Novel miRNAs continue to be discovered throughout animals and plants. More and more miRNAs are found to be non-conserved and usually express in specific species, tissues or cells at a relatively low level. Identifying and studying these novel specific expressed miRNAs will help us to better understand the fine regulation of the cell and life. However, annotating a small RNA as a *bona fide* miRNA should be cautious to avoid false identification because deep sequencing technology has revealed a massive amount of small RNAs with somewhat similar characteristics to miRNAs. Recently, Fromm *et al.* updated a uniform system for the annotation of miRNAs [Fromm et al., 2015]. To avoid mis-annotation, we strictly followed these latest updated miRNA annotation criteria. First, the reads of the nine novel miRNAs were at least more than the median value of all known miRNAs in this deep sequencing dataset. Because deep sequencing data are considered as good evidence for gene expression, the criterion of the confident expression of the newly identified miRNAs was well met in this study. Second, all miRNAs, except for miR-novel-16, started with 5′-end U or A, ranged from 20-24 nt. Third, all miRNA reads for each miRNA had 100% 5′-end homogeneity. Forth, all miRNAs have the typical hairpin characteristics satisfied with the miRNA annotation criteria. For example, the stem region has at least 16-nt complementary base-pairing; the loop sequence is 8-40 nt in length. Fromm *et al*. also stated that there should be two 20–26 nt long reads expressed from each of the two arms of the hairpin precursor with 2-nt 3′ overhangs. However, the ratio of mature miRNA/miRNA* is usually very asymmetric with a discrepancy of >10,000:1 [Liu et al., 2008]. Therefore, it is very hard to get reads for the less abundant miRNA* when the mature miRNA expressed at a low level. Especially, many species- or cell-specific and evolutionally young miRNAs usually expressed at relatively low levels. For this reason, although this criterion is evidence of Drosha/Dicer processing, it was not applied in this study. However, we did identify the miRNA* reads of two novel miRNAs, miR-novel-27 and miR-novel-65, with 2-nt 3′ overhangs. In addition, the conservation criterion was not applied too because many non-conserved miRNAs have been robustly validated. In fact, we even found that miR-novel-27 is unique to the mouse species and originated from a recent inverted local half hairpin duplication event. In this case, there is no conservation between species at all. Overall, these nine novel miRNAs are well satisfied with the recently updated miRNA annotation criteria and are backed robust supporting of *bona fide* miRNAs.

MiRNAs are believed to rapidly evolve and show high frequent birth and death in the course of evolution [Meunier et al., 2013; Zhang et al., 2018]. *De novo* miRNA emergence started from the formation of a pre-miRNA-like hairpin structure that was eventually to be recognized by Drosha/Dicer. In plants, hairpins commonly generated by inverted duplication of target genes [Allen et al., 2004; Axtell et al., 2011]. In animals, hairpin generally evolved from the initially unstructured sequence in random [Axtell et al., 2011; Zhang et al., 2018]. Another major source for the emergence of new miRNA genes in animals was from local duplication of existing miRNA genes (full hairpin), followed by subfunctionalization and neofunctionalization [Zhang et al., 2018]. In this way, miRNA genes expanded to gene clusters to form miRNA families [Zhang et al., 2018]. But, the second mechanism only explains the miRNA expansion and evolvement from an existing miRNA gene but not the *de novo* birth of the existing miRNA gene itself. So, this mechanism is actually a miRNA family expansion mechanism but not the novel miRNA *de novo* emergence mechanism. Are there other sources of the *de novo* miRNA emergence other than from unstructured sequences? Here, we represented a new type of *de novo* miRNA birth mechanism exampled by the miR-novel-27. This new mechanism is the first example of inverted duplicating a short genomic sequence and form a palindromic sequence in animals, from which an RNA molecular could fold into a full hairpin structure. We called it “inverted local half hairpin duplication” in order to distinguish from the previously demonstrated local duplication of existing miRNA genes (full hairpin). Overall, this study presented a new model of *de novo* miRNA gene birth mechanism in animals supported by miR-novel-27.

## Acknowledgment

This work was supported by National Natural Science Funds for Distinguished Young Scholar (31100570) and the Science and Technology Supporting Program of Jiangsu Province (BE2013657). This work was also supported by the Central Laboratory for Translational Medicine of Qidong People’s Hospital.

## Supplemental Figure Legends

**Figure S1.**
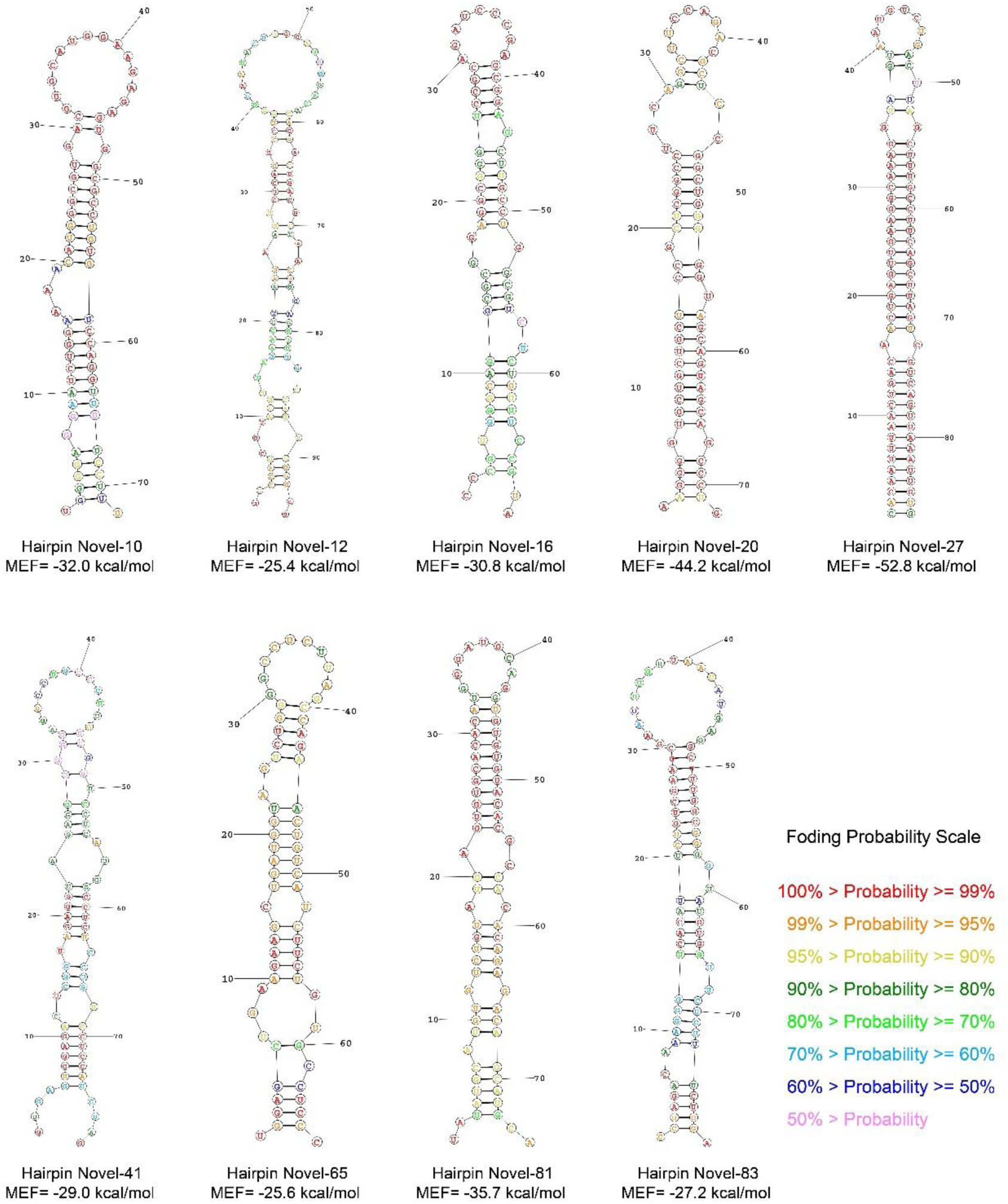
The folded structures of the candidate miRNA hairpins. The colors of nucleotides showed the base-pairing probabilities as indicated.

**Figure S2.**
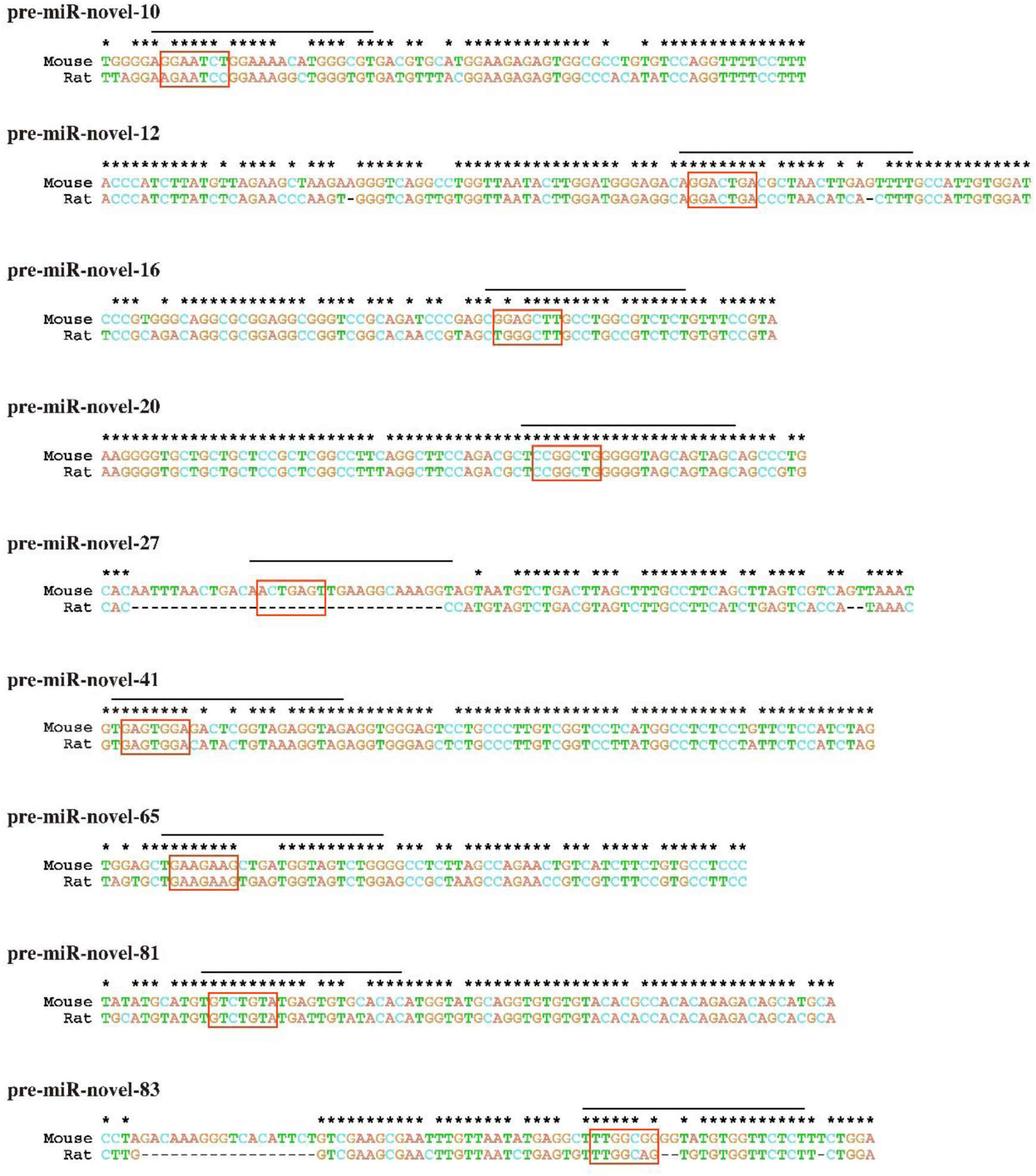
The sequence alignments of the candidate precursors between mouse and rat species. Mature miRNA sequences were indicated with lines at the top, and seed regions were labeled by red boxes.

**Figure S3.**
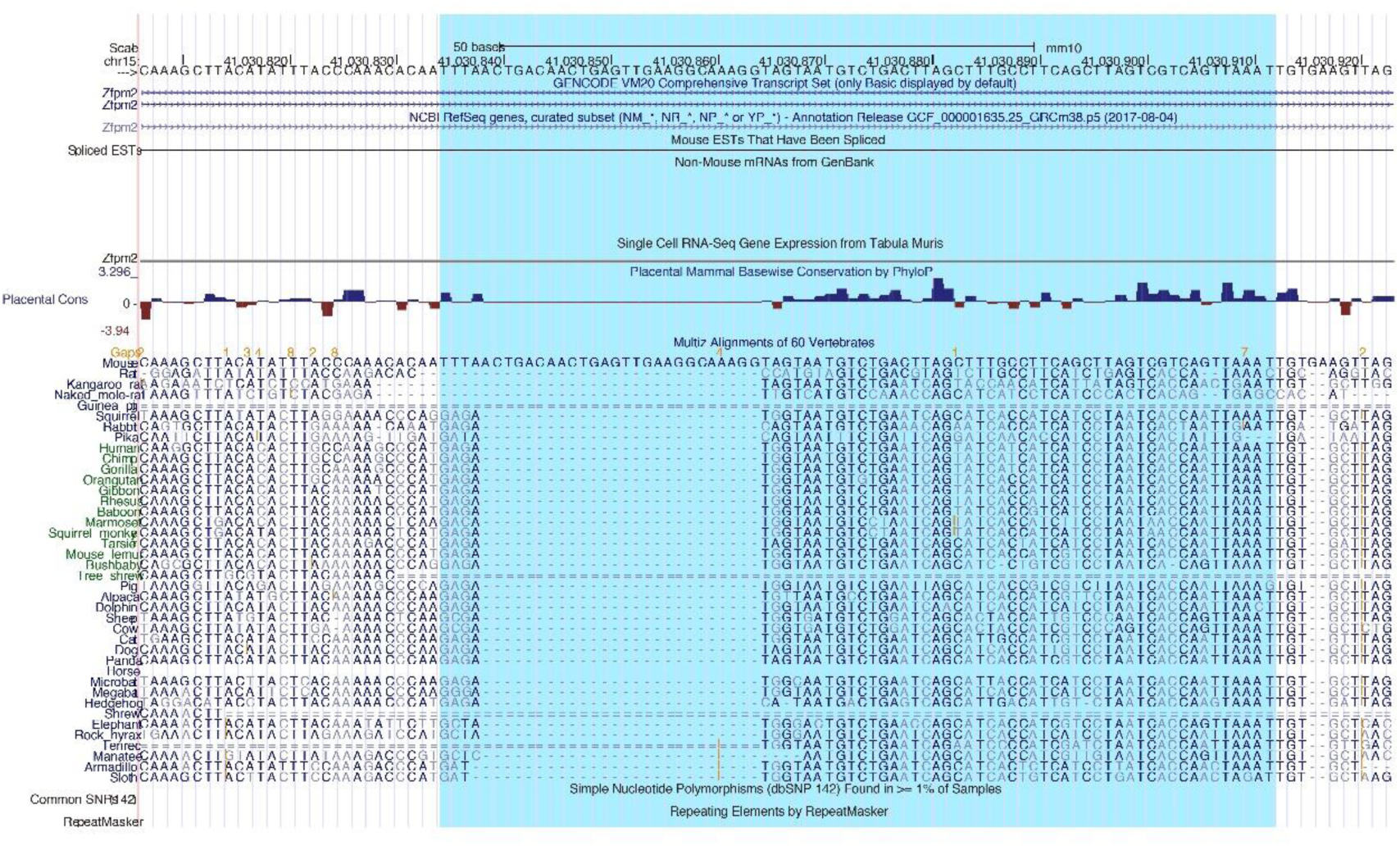
MiR-novel-27 is unique to mouse species. the surrounding sequence of miR-novel-27 was aligned to other *Placentalia* animals. The blue shadow indicated the hairpin sequence region. The image was extracted from the UCSC Genome Browser.

**Figure S4.**
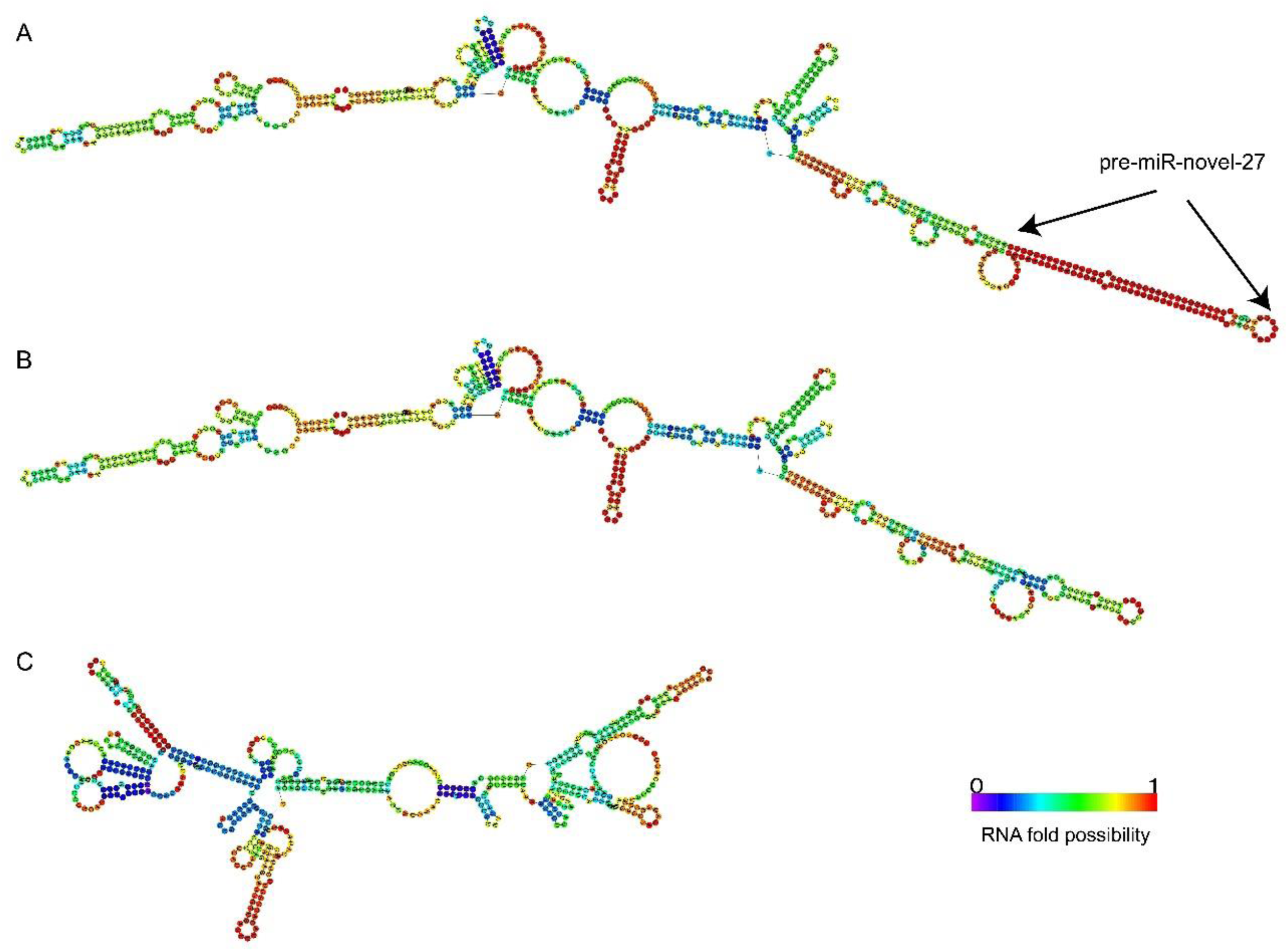
Folded structures of the miR-novel-27 surrounding sequence. (A) The miR-novel-27 precursor sequence was extracted along with the +/-300-nt surrounding sequence for RNA structure folding analysis. Pre-miR-novel-27 was stably formed within the indicated context sequence. (B) The RNA sequence use in A was folded again after deleting the duplicated 5′ arm region. No pre-miRNA-like hairpin formed throughout the RNA molecular. (C) Sequence from the according region of the rat species was folded and no pre-miRNA-like hairpin formed. Fold probability was scaled as indicated.

